# Adaptation of reach action to a novel force-field is not predicted by acuity of dynamic proprioception in either older or younger adults

**DOI:** 10.1101/2020.07.13.200733

**Authors:** Nick M. Kitchen, R Chris Miall

**Affiliations:** School of Psychology, University of Birmingham, Birmingham, UK; Dept. of Speech & Hearing Science, University of Washington, Seattle, WA, USA

**Keywords:** Proprioception, Ageing, Reaching, Force-Field Adaptation

## Abstract

Healthy ageing involves degeneration of the neuromuscular system which impacts movement control and proprioception. Yet the relationship between these sensory and motor deficits in upper limb reaching has not been examined in detail. Recently, we reported that age-related proprioceptive deficits were unrelated to accuracy in rapid arm movements, but whether this applied in motor tasks more heavily dependent on proprioceptive feedback was not clear. To address this, we have tested groups of younger and older adults on a force-field adaptation task under either full or limited visual feedback conditions and examined how performance related to dynamic proprioceptive acuity. Adaptive performance was similar between the age groups, regardless of visual feedback condition, although older adults showed increased after-effects. Physically inactive individuals made larger systematic (but not variable) proprioceptive errors, irrespective of age. However, dynamic proprioceptive acuity was unrelated to adaptation and there was no consistent evidence of proprioceptive recalibration with adaptation to the force-field for any group. Finally, in spite of clear age-dependent loss of spatial working memory capacity, we found no relationship between memory capacity and adaptive performance or proprioceptive acuity. Thus, non-clinical levels of deficit in dynamic proprioception, due to age or physical inactivity, do not affect force-field adaptation, even under conditions of limited visual feedback that might require greater proprioceptive control.

## Introduction

Typical healthy ageing brings about a number of notable changes to the neuromuscular system which influences the ability to control simple movements. This includes loss and remodelling of musculature and motor units (Lexell, 1995; Morley et al., 2001; Slack et al., 1979), as well as degeneration of the peripheral nerves and neuromuscular junction (Jacobs & Love, 1985; Valdez et al., 2010). Collectively, these changes contribute to the characteristic increases in variability, duration and online corrections that are seen in basic upper limb movements of older adults (Contreras-Vidal et al., 1998; Darling et al., 1989; Helsen et al., 2016; Ketcham et al., 2002; Kitchen & Miall, 2019; Seidler et al., 2002; Yan et al., 2000). Some of these kinematic features are also thought to play a compensatory role to preserve endpoint accuracy for older adults during targeted reaching tasks (Helsen et al., 2016; Kitchen & Miall, 2019; Lee et al., 2007; Seidler-Dobrin & Stelmach, 1998).

Ageing also involves degeneration of proprioceptive sensory organs in the neuromuscular system responsible for sensations of limb position and motion (Proske & Gandevia, 2012). Human and animal studies have shown age-related degeneration of the muscle spindle, including a reduced number of intrafusal fibres (Kararizou et al., 2005; Swash & Fox, 1972), increased capsular thickness (Swash & Fox, 1972) and degraded morphology of primary spindle endings (Kim et al., 2007) that place limits on the perceptual acuity of proprioceptive sensations in advanced age. Indeed, a broad range of proprioceptive assessment techniques – both passive and active in nature – have shown age-dependent loss of proprioceptive acuity (Adamo et al., 2007; Cressman et al., 2010; Herter et al., 2014; Lei & Wang, 2018; Schaap et al., 2015; for review see Goble et al., 2009), and acceleration of loss with physical inactivity (Adamo et al., 2009; Helsen et al., 2016; Kitchen & Miall, 2019; Wright et al., 2011). Given that complete loss of proprioceptive sensation has profoundly detrimental effects on upper limb movements (Miall et al., 2018; Sarlegna et al., 2010; Yousif et al., 2015) it is clear that controlled motor performance is dependent on these sensory signals. Yet in spite of reports showing age-dependent proprioceptive decline and age-dependent reduction in motor function in the lower limb (Hurley et al., 1998; Lord et al., 1991; Sorock & Labiner, 1992; Wingert et al., 2014) only recently has there been increased interest in this issue for the upper limb.

Helsen et al. (2016) investigated this sensorimotor relationship in the wrist using two proprioceptive assessment tasks (passive movement detection and ipsilateral position matching) and a rapid movement task to visual targets. In line with other reports (Adamo et al., 2009; Kitchen & Miall, 2019; Wright et al., 2011), physically inactive older adults were found to have reduced proprioceptive acuity, with increased movement detection thresholds and position matching errors. They also made more online corrective adjustments during movements, though endpoint accuracy was relatively unaffected. Critically, no relationships were found between proprioceptive acuity and either aiming errors or movement kinematics. However, in order to understand this relationship in a more naturalistic or real-world setting, the proprioceptive tests should be active, to incorporate sense of effort and motor efference which can influence position sense (‘t Hart & Henriques, 2016; Smith et al., 2009). Active testing can even reduce position matching errors in both older and younger adults (Erickson & Karduna, 2012; Langan, 2014; Lönn et al., 2000). We therefore tested groups of younger and older adults on two tasks that separately assessed active proprioceptive acuity and rapid target reaching performance (Kitchen & Miall, 2019). Like Helsen et al. (2016), we found age effects on motor performance. We also found that physically inactive older adults had increased systematic (but not variable) proprioceptive errors, but there were no associations between proprioceptive acuity and motor errors for either age group (Kitchen & Miall, 2019).

However, those tasks emphasised speed, where it seems likely that sensory feedback control is minimized in favour of predictive, feedforward processes (Miall & Wolpert, 1996; Wolpert et al., 1995). Thus, the question remains as to how sensory impairments might influence performance on tasks which require greater sensory feedback control (for review see Shadmehr et al., 2010). To perturb proprioceptive feedback specifically, external forces can be applied during discrete targeted movements (Krakauer et al., 1999; Sarlegna et al., 2010; Shadmehr & Mussa-Ivaldi, 1994). Unlike tasks with visual feedback perturbation (Anguera et al., 2011; Bock, 2005; Buch et al., 2003; Contreras-Vidal et al., 2002; Hegele & Heuer, 2010; Seidler, 2006; Vandevoorde & Orban de Xivry, 2019), a number of studies have indicated a minimal role of ageing in the ability to adapt movements to novel force environments (Cesqui et al., 2008; Huang & Ahmed, 2014; Rajeshkumar & Trewartha, 2019; Reuter et al., 2018; Trewartha et al., 2014). Nonetheless, several age-dependent performance predictors have been highlighted, including increased muscle co-contraction (Huang & Ahmed, 2014), reduced central processing of kinematic errors (Reuter et al., 2018) and reduced spatial working memory capacity (Trewartha et al., 2014). Furthermore, younger adults demonstrate a tight relationship between proprioceptive perception and motor performance when adapting to novel force-fields (Haith et al., 2008; Mattar et al., 2013; Ohashi et al., 2019; Ostry et al., 2010). Moreover, whilst adaptation to forces is possible in younger adults without visual feedback of hand position (Franklin et al., 2007; Lefumat et al., 2015; Scheidt et al., 2005), all previous studies with older adults have included either full (Cesqui et al., 2008; Huang & Ahmed, 2014; Rajeshkumar & Trewartha, 2019; Trewartha et al., 2014) or early visual feedback (first half of the movement; Reuter et al., 2018). Hence, the interactions of age and proprioceptive acuity on dynamic motor adaptation remains poorly understood, and the contributions of visual and proprioceptive feedback to motor adaptation have not been dissociated.

We have therefore investigated the relationship between dynamic proprioception and force-field adaptation in older and younger adults. Participants were given either full or limited visual feedback during force-field adaptation, and dynamic proprioceptive assessments were made before and at intervals throughout the adaptation task. Spatial working memory capacity and physical activity status were also examined, collectively providing a comprehensive overview of reach adaptation to novel field dynamics and its interaction with proprioceptive acuity across the lifespan.

## Methods

### Participants

A total of 34 older (13 male, 75.6 ± 7.2 yrs) and 35 younger (3 male, 19.1 ± 0.9 yrs) adults participated in the experiment, all of which were right-handed (30 or higher on the 10-item Edinburgh Handedness Inventory; Oldfield, 1971). Participants were excluded from the study for history of neurological illness, carpal tunnel syndrome, arthritis or similar movement pains or limitations in the arm, wrist or fingers. Participants were also screened to ensure no prior experience with adaptation to novel field dynamics. Older adults completed the Montreal Cognitive Assessment (MoCA) and were only included in the analysis if they scored 26 or above out of 30, which is indicative of normal cognitive functioning (Nasreddine et al., 2005). This study was approved by the University of Birmingham ethics panel and all participants gave written consent prior to experiment performance.

Older and younger participants were randomly allocated to two subgroups (see section A(ii) *Visual feedback conditions*); 17 older adults in Vision+ (OAVis+; 5 male, 75.2 ± 8.3 yrs), 17 older adults in Vision-(OAVis-; 8 male, 75.9 ± 6.2 yrs), 17 younger adults in Vision+ (YAVis+; 1 male, 18.9 ± 0.8 yrs) and 18 younger adults in Vision-(YAVis-; 2 male, 19.4 ± 1.0 yrs). The visual feedback sub-groups were well matched for age in both older (t[32] = −0.28, *p* = .780) and younger (t[33] = −1.69, *p* = 0.101) adult groups.

### Physical Activity

Self-reported physical activity levels were measured using the Physical Activity Scale for the Elderly (PASE; Washburn et al., 1993) and the IPAQ-Short questionnaire (Craig et al., 2003) for older and younger participants respectively. Based on a median score threshold of 155.4 from the PASE questionnaire, older adults were assigned to either a physically active (n = 17, 219.4 ± 61.9) or inactive (n = 17, 108.7 ± 40.7) group. Likewise, a median score of 1988 MET-min per week on the IPAQ was used to sub-divide younger adults into physically active (n = 17, 3763.9 ± 1253.2 MET-min per week) and inactive (n = 18, 1082.7 ± 531.1 MET-min per week) groups. These sub-groups were used to examine interactions of physical activity and proprioception.

### Spatial Working Memory Assessment

Prior to completing the experiment, spatial working memory capacity was assessed using a modified search task previously used with older adults and cognitively impaired patient populations (Kessels et al., 2010; Van Asselen et al., 2005). Participants were presented with sets of 4-10 blue tiles on a computer monitor. They were instructed to search for one hidden target, by selecting tiles one at a time. They continued this until they chose the target tile, briefly revealed as a green tick symbol before reverting back to blue (a sequence we term a “search”). Participants were required to remember this tile location and then continue searching for a new target, which might appear on any of the remaining tiles. They had to complete searching until all possible targets had been found (i.e. as many searches as there were tiles). In any given search, if the participant chose a tile which they had previously selected in that search, a red cross symbol briefly appeared; this was defined as a within-search error. Similarly, if participants selected a tile which had been the target in an earlier search, they also saw a red cross; this was a between-search error. The purpose of the task was therefore to find all of the green ticks whilst minimising the number of search errors. All participants completed 2 trials at each of 4, 6, 8 and 10 tiles after an initial 2 trials with 4 tiles as a familiarisation. To minimise distractions, participants also wore headphones that played distinct audio tones associated with selection of a blue tile, finding a target, or making an error. Between and within search errors were averaged across the 2 attempts at each number of tiles, then compared between age groups. After reviewing the data, we found that only a single error was made across all participants in the 4 tile condition, so it was not included in the main analysis; this left the 6, 8 and 10 tile conditions. Total search errors (the sum of between- and within-search errors across these three tile conditions) were also used as a correlate for performance in the other tasks.

### Main Experimental Set-Up

The main tasks were performed on a low-inertia and near-frictionless 2D-planar manipulandum (vBOT; Howard, Ingram, & Wolpert, 2009) which made it possible to record reaching movements in a 40×64cm workspace (Figure 1A). Participants placed their forehead on a padded headrest and looked down onto a large, horizontally mounted, mirrored surface that reflected targets and visual feedback images from an LCD screen directly above. Participants grasped the manipulandum handle with their right hand underneath the mirrored surface, which blocked any direct vision of the arm or hand. The start position and target were displayed as white and grey 1cm radius markers respectively, with the handle position displayed as a white 0.5cm radius marker, when feedback was available. Start position was located in the midline, 8cm into the workspace (roughly 28cm from participant’s chest) with the target located 20cm directly ahead along the mid-sagittal axis. The start position and target remained in the same position and were kept visible at all times during task performance.

**Figure 1.**
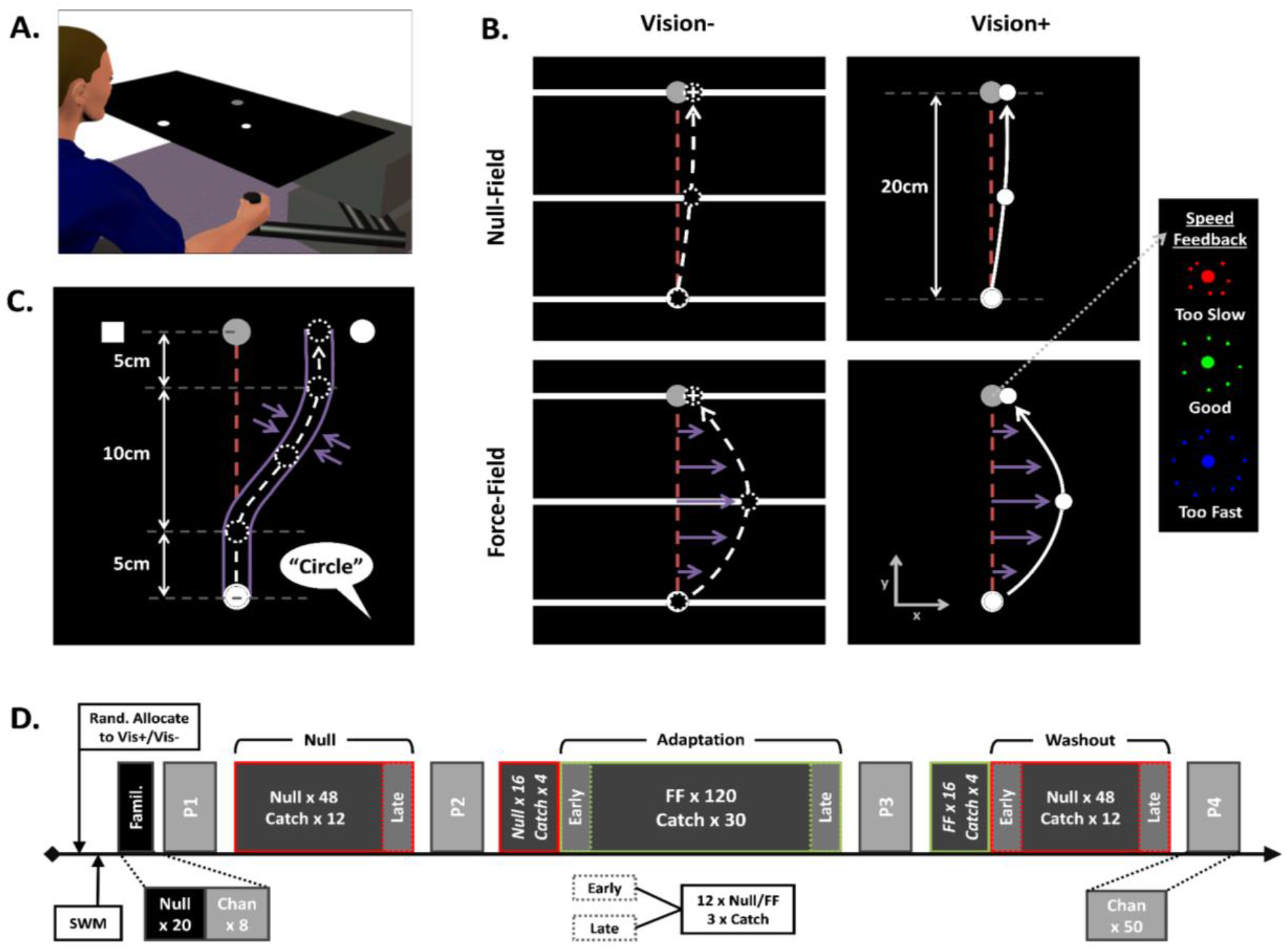
Experimental set-up, tasks and design. **A.** vBOT handle underneath reflective surface that displayed images during task performance. **B.** Force-field task and visual feedback conditions. The Vision-group (left column) were shown a horizontal white bar as a hand position cursor (which provided movement feedback only along y-axis regarding distance to target), as well as white cross terminal position feedback. The Vision+ group (right column) were provided with full visual feedback (white circular cursor) throughout. Purple arrows show forces imposed by velocity-dependent field (lower row), with furthest right panel illustrating feedback provided at the target, after trial termination, on movement speed **C.** Dynamic proprioception task. Participants actively moved through a minimum jerk pathway constrained by stiff virtual walls before verbally indicating which side of the target they felt they had been guided to (“Square” or “Circle”). **D.** Experimental design. Participants were randomly allocated to either the Vision+ or Vision-group before completing the spatial working memory task (SWM) and brief familiarisation task. Blocks (P1-P4) of channel trials (Chan) for proprioceptive assessment were interleaved between target reaching blocks of null-field (red outlined boxes) and force-field (green outlined boxes) trials. Data was averaged across early and late portions of blocks for statistical analyses.

### Task A. Force-Field Adaptation

#### (i) Procedure

A summary of this task is displayed in Figure 1B. To begin, participants first moved to the start position, which turned orange, and then made a reaching movement towards the target. Once they had moved the required 20cm ahead, they intersected a soft virtual wall which ran orthogonally through the target. Regardless of whether the target was hit or not, it then displayed an ‘explosion’ graphic that provided participants with colour-coded feedback regarding the speed of their movement compared to a desired movement duration of 350-450ms (Figure 1B). The target then turned grey again and participants were actively guided back to the start position by a spring force (500 N.m^−1^, 1 N.m.s^−1^ damping) for the next trial.

Movements during null-field trials were unconstrained. However, force-field trials were performed in the presence of a velocity-dependent force-field applied to the vBOT handle as described by Equation 1.

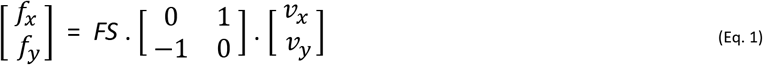

Where *f*_*x*_ and *f*_*y*_ refer to the imposed forces on the vBOT handle, *FS* refers to the field strength which was held constant at 15N.m^−1^.s^−1^, and *v*_*x*_ and *v*_*y*_ refer to the velocity of the vBOT handle. This “curl-field” (Shadmehr & Mussa-Ivaldi, 1994) perturbed movements in a clockwise direction that deviated movements from their normally linear path towards the target (Figure 1B).

Catch trials were presented randomly within the final 3 of every set of 5 null- and force-field trials. In these trials, hand position was constrained to a linear path between start position and target using stiff virtual walls (stiffness: 2000 N.m^−1^ with 10 N.m.s^−1^ damping imposed by vBOT motors). Forces against the channel wall were used as a measure of participants’ compensation for the force-field during the course of adaptation. Visual feedback of distance without direction was presented as a white semi-circular arc whose radius increased proportionally to the linear distance travelled towards the target (see Lago-Rodriguez & Miall, 2016). Coloured speed feedback and active guidance back to start were still provided for these trials.

Participants were informed that they may experience some forces at the vBOT handle during target reaching movements, but that their objective was to try and keep movements as straight and as accurate as possible (keeping the white cross on target for Vision- and hand position cursor on target for Vision+), whilst also trying to maintain appropriate movement speed (green explosions).

#### (ii) Visual Feedback Conditions

Participants were given either full visual feedback (Vision+) or limited visual feedback (Vision-) during the experiment. For the Vision+ sub-group, the white circular hand position cursor was available at all times. The Vision-group were shown a moving horizontal white bar as a hand position cursor, which only provided visual feedback regarding movement distance to the target (Figure 1B, left column). This meant that any lateral movements (and hence features of the perturbation) were hidden from view at all times. To reduce chances of proprioceptive drift occurring (Brown et al., 2003a, 2003b), terminal position feedback (the location at which participant intersected the endpoint orthogonal soft wall) was also provided at the end of each movement (1cm width white cross). Movement speed feedback (coloured explosion) and active guidance back to the start position after trial termination were identical for Vision+ and Vision-sub-groups. Visual feedback during catch trials was identical, regardless of sub-group (see A(i) above).

#### (iii) Outcome Measures and Analysis

Lateral hand deviation from the linear path connecting start position and target was recorded at peak hand velocity (peak-velocity lateral deviation; PVLD). For catch trials, the force generated against the channel wall was compared with the ideal force necessary to perfectly compensate for the force-field given the velocity profile of the movement. Specifically, we fitted a least-squared linear regression model (without an intercept) to the observed versus optimal orthogonal forces for each trial and used the slope as an adaptation index (Huang & Ahmed, 2014; Trewartha et al., 2014); a value of +1 indicated complete compensation for the force-field.

To examine kinematic performance, we recorded peak hand velocity, total movement duration and the time to peak velocity (expressed as a percentage of total movement duration). Movement duration was defined as the time taken to move from 1cm beyond start position to 1cm before target. Trials where total movement duration was greater than 1.5 seconds were not included in the analysis (this equated to < 1 % of trials).

To monitor performance change across the experiment, adaptation and kinematic measures were averaged across the first and last 15 trials (3 catch trials, and 12 null/force-field) in a given block which we term “early” or “late” respectively (this excluded the additional 20 null and 20 force-field trials at the start of the adaptation and washout blocks respectively; see Figure 1D for details). Extent of adaptation was then defined as the positive difference between early and late performance in the adaptation block: reduction in lateral error (early minus late PVLD) and increase in adaptation index (late minus early).

### Task B. Dynamic Proprioception

#### (i) Procedure

The details of this task have been described previously (Kitchen & Miall, 2019) and are illustrated in Figure 1C. Participants made reaching movements towards the target that were tightly constrained to a pre-determined trajectory using stiff virtual walls (stiffness: 2000 N.m^−1^ with 10 N.m.s^−1^ damping imposed by vBOT motors). This channel deviated the hand path laterally from the target through a minimum jerk profile. No forces were applied in the forward direction; no visual feedback was provided. At the end of the movement, a white square and circle appeared at constant positions on the left and right-hand side of the target respectively. The participant then made a verbal 2AFC response (“Square” or “Circle”) to indicate the side of the target that they felt they had been guided to. Following their verbal response, participants were actively guided back to the start position by a spring force (500 N.m^−1^, 1 N.m.s^−1^ damping) before starting the next trial. The size and direction of the lateral deviation was manipulated on a trial-by-trial by two adaptive staircase sequences.

#### (ii) PEST Sequences

The lateral deviation imposed by the virtual channel walls was defined by 2 randomly interleaved PEST sequences (Taylor & Creelman, 1967), with one starting from the left side of the target (“Square”) and the other from the right (“Circle”). This was designed to minimize awareness of sequence progression and improve convergence of lateral deviations towards participants true perceptual thresholds. Each sequence started at a deviation magnitude of 3cm (±0.05cm added noise) with an initial step size of ±1cm. This step either increased or decreased the deviation magnitude (or “level”) according to the cumulative accuracy of the verbal responses from the two repeats which were performed at each level of the sequence. If both responses were correct, then the deviation magnitude would decrease, and if both were incorrect it would increase. When there was a tie (1 correct and 1 incorrect response) a third repeat at that level was performed, with the subsequent response determining the overall accuracy of the responses to that level. Whenever the sequence reversed direction (a successive step increase and decrease in deviation magnitude or vice-verse) the new step size became half of the previous one i.e. from 1cm to 0.5cm at the first reversal. The interleaved PEST sequences continued in this manner for 50 trials, which constituted one proprioceptive assessment block (Figure 1D).

#### (iii) Outcome Measures and Analysis

The verbal responses indicating perceived limb position were converted to binary values (“Circle” = 1, “Square” = 0) and expressed as a proportional response probability for each level of the deviation before being fitted with a logistic function using the Matlab function glmfit. To remove the effects of outlying response values on curve fitting, data points with a Pearson residual that was greater than 2.5 standard deviations away from residual mean were excluded from analysis (this equated to <1% of data). From the logistic we then estimated the bias and the uncertainty range as indices of proprioceptive acuity. The bias represents a systematic error in limb perception and corresponds to the 50^th^ percentile of the logistic function; a positive bias here represents a perception of hand position to be further to the right (“Circle”) of the target and negative bias indicates a perceptual error to the left (“Square”). The uncertainty range gives a variable error of perceived hand position and is given as the interval between the 25^th^ and 75^th^ percentile of the logistic function.

In addition to verbal perceptual responses, the average movement velocity was calculated for the portion of the movement from 1cm beyond the start position to 1cm before the target. The average orthogonal forces exerted on the virtual channel walls were also recorded in the middle of the final, straight portion of the movement (16-19cm of the 0-20cm movement). These measures were used as correlates for proprioceptive measures to ensure minimal confounding effects of effort on perceptual judgements of limb position (Smith et al., 2009).

### Experimental Design

The full design is shown in Figure 1D. The experiment started with a familiarisation of 20 null-field trials (self-selected speed with full hand position cursor feedback regardless of visual feedback condition) and 8 proprioceptive channel trials. Participants then performed the first block of 50 proprioception trials (P1), followed by a block of 60 target reaching movements trials (48 null-field and 12 catch; “Null”), before another proprioception block (P2). The main block of target reaching trials was then performed, consisting of 16 null-field trials (interleaved with 4 catch trials) and 120 force-field trials (interleaved with 30 catch trials; “Adaptation”). This was immediately followed by the third proprioceptive block (P3) and a final block of target reaching trials consisting of 16 force-field trials (interleaved with 4 catch trials) and 48 null-field trials (interleaved with 12 catch trials; “Washout”). The fourth and final proprioceptive block (P4) was then performed to finish.

### Statistical Analysis

Both proprioceptive and adaptation data were normalised to baseline performance (block P1 for proprioception, late Null data for adaptation; Figure 1D) before analysis with mixed-design ANOVAs that included between-subjects factors of age group (older or younger) and visual feedback condition (Vision+ or Vision-) and a within-subjects factors of time-point (early or late in the block for adaptation data; test blocks P2, P3, P4 for proprioceptive data). Adaptation and washout blocks were analysed separately to examine finer aspects of group performance for these respective stages of the experiment. To probe the relationship between proprioception and adaptation, adaptation extent was separately correlated with baseline proprioceptive acuity and with change in bias after the adaptation (P2 subtracted from P3).

Since the baseline (P1) proprioceptive assessment was made prior to exposure to the visual feedback condition, these data were separately analysed in two-way ANOVAs that included only age group and physical activity status (active or inactive) as between-subjects factors. Physical activity indices were converted to z-scores within the two age groups, and used as a predictor of adaptation extent in multiple linear regression models that controlled for age and visual feedback condition grouping. Finally, spatial working memory scores were analysed in mixed ANOVAs with a between-subjects factor of age group and within-subjects factor of memory load (6, 8 or 10 tiles).

In an attempt to meet parametric assumptions whilst minimizing data exclusion, we examined several approaches to deal with outlying data. Namely, we compared the number of significant Shapiro-Wilk tests reported for all subsets of data included in our statistical analyses, testing the raw data, log-transformed data, and data with outliers removed according to 2 different thresholds (3 and 2.5 standard deviations [SD] from the group mean at each repeated measure level). Whilst the log transformation and the 3 SD threshold outlier removal did reduce the number of significant Shapiro-Wilk tests compared to raw data, there were still a concerningly high number of non-normal distributions from these approaches. Conversely, the 2.5 SD threshold removal minimized the number of non-normal distributions with relatively low data exclusion overall (4.0% of all data), so we selected this approach for our analyses. All values are presented as group means ± standard deviation unless otherwise stated and in all cases where the sphericity assumption was violated a Greenhouse-Geisser correction was used, with significance assessed at the *p* < .05 level. In cases of more than two multiple comparisons (post-hoc tests, correlations and regression models) we report *p*-values which have been adjusted using a false discovery rate analysis (Benjamini & Hochberg, 1995; Yekutieli & Benjamini, 1999; also see use in Vandevoorde & Orban de Xivry, 2019), denoted as *p*_*adj*_. The Bayes factor (*BF*_*01*_) associated with correlations between measures from different tasks was also calculated using the program JASP (JASP Team, 2020) to provide further evidence for (or against) non-significant Pearson’s correlations (Wagenmakers, Love, et al., 2018; Wagenmakers, Marsman, et al., 2018). All correlations were calculated as two-tailed tests using a uniform prior distribution. Values for *BF*_*01*_ ranged from 0.51 (anecdotal evidence for the alternative hypothesis) to 4.49 (moderate evidence for the null hypothesis), though some were also close to 1 indicating equivocal evidence for both (Wagenmakers, Love, et al., 2018).

## Results

### Force-Field Adaptation

#### Lateral deviation

Mean lateral deviation at peak velocity (PVLD) across the entire experiment, before outliers were removed, are shown in Figure 2A (Vision+ groups) and 2B (Vision-groups), with group PVLD (normalized to baseline) for early and late portions of the adaptation and washout block shown in Figure 2C and 2D, respectively. Overall, participants made expected reductions in PVLD from early to late in the adaptation block (F[1, 60] = 159.36, p < 0.001, *η*^*2*^_*p*_ = 0.73). There was a timepoint x visual feedback interaction on PVLD (F[1, 60] = 4.34, p = 0.041, *η*^*2*^_*p*_ = 0.07), though post-hoc tests revealed no difference between the Vis+ and Vis-groups at either timepoint (both p ≥ 0.123). The timepoint x age group interaction was also significant (F[1, 60] = 5.29, p = 0.025, *η*^*2*^_*p*_ = 0.08), where younger adults made larger errors (3.2 ± 1.3cm) than older adults (2.4 ± 1.1cm) early in the adaptation block (t[62] = −2.64, p = 0.010; Figure 2C). The timepoint x age group x visual feedback interaction was not significant (p = 0.537) and there were no further effects or interactions of age and visual feedback on PVLD (all p ≥ 0.073).

**Figure 2.**
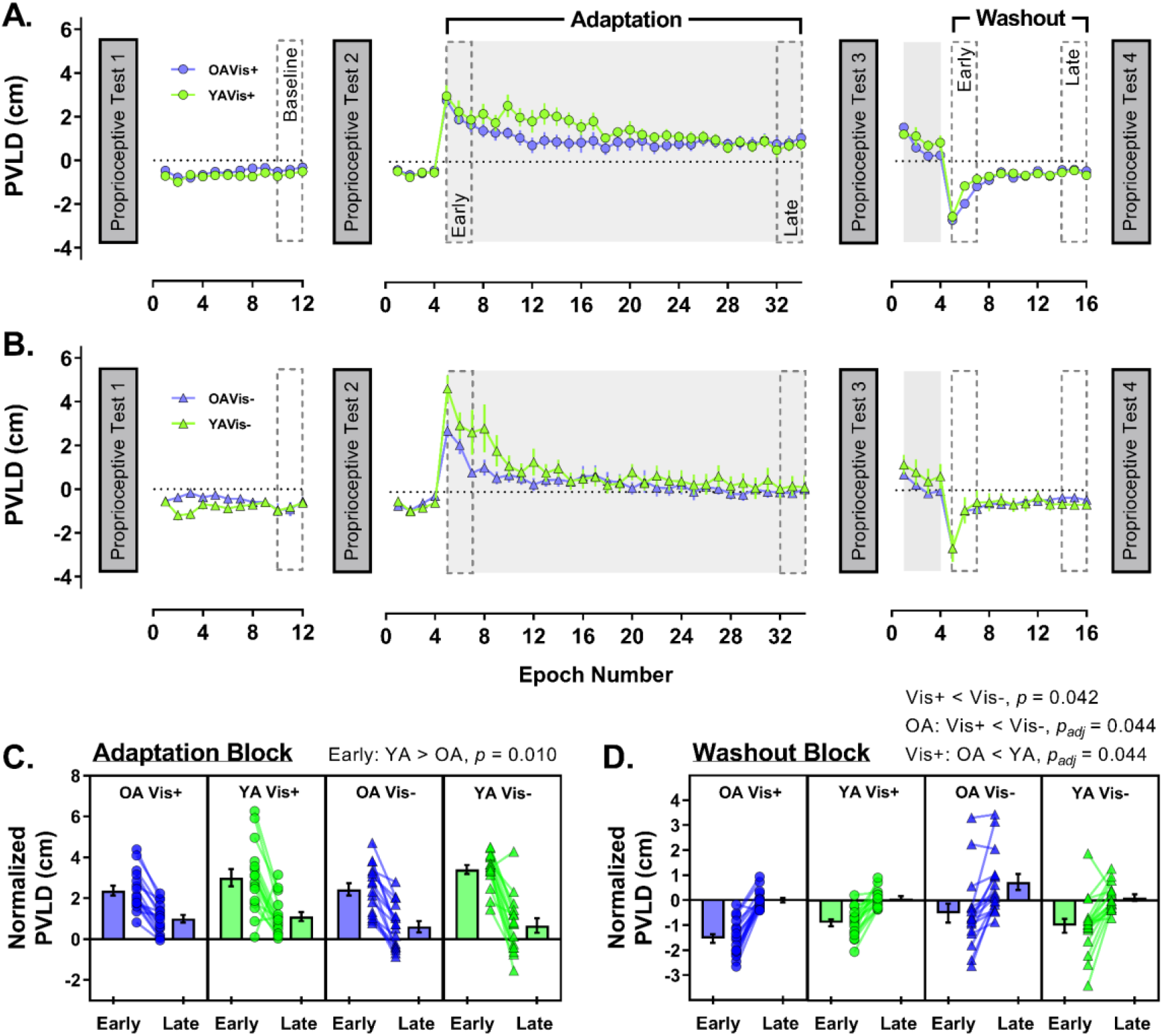
Group peak velocity lateral deviation (PVLD) data across the experiment for older (purple) and younger (green) adults in (**A**) Vision+ condition (circles) and (**B**) Vision-condition (triangles). For visualization purposes only, epochs of 4 trials were created for each participant and then averaged within groups to generate the group means (± 1SE) shown in panels **A** and **B**, with grey shaded regions representing periods where the force-field was on. Lower panels show PVLD data (normalized to baseline) for the early and late portions of the adaptation block (**C**) and the washout block (**D**). In these panels, the means of individual participants are shown as symbols, with group averages (± 1SE) displayed in neighbouring bars. Significant post-hoc comparisons for group interactions also shown above respective panels.

PVLD was reduced between early and late washout (F[1, 63] = 138.51, p < 0.001, *η*^*2*^_*p*_ = 0.69), but there were no timepoint interactions (all p ≥ 0.136). The Vis+ group (−0.59 ± 0.46 cm) showed slightly increased movement errors than the Vis-group (−0.19 ± 1.06 cm) across the whole washout block (F[1, 63] = 4.30, p = 0.042, *η*^*2*^_*p*_ = 0.06). The age x visual feedback interaction was significant (F[1, 63] = 5.40, p = 0.023, *η*^*2*^_*p*_ = 0.08), with post-hoc tests indicating that for older adults only, the Vis+ group (−0.77 ± 0.47cm) made larger PVLD errors than the Vis-group (0.09 ± 1.3 cm; t[18.7] = −2.49, *p*_adj_ = 0.044; Figure 2D). Furthermore, for the Vis+ group only, older adults (−0.77 ± 0.47 cm) made larger PVLD errors than younger adults (−0.40 ± 0.38 cm; t[32] = −2.46, *p*_adj_ = 0.044; Figure 2D). The remaining post-hoc comparisons were not significant (all *p*_adj_ ≥ 0.197). There was no effect of age group on washout PVLD either (*p* = 0.651).

#### Channel trials

Adaptation index values across the experiment (before outlier exclusion) are shown in Figure 3A (Vision+ groups) and 3B (Vision- groups), with group adaptation index (normalized to baseline) for early and late portions of the adaptation and washout block shown in Figure 3C and 3D, respectively. Channel trial data showed that adaptation index increased from early to late adaptation as expected (F[1, 61] = 75.09, p < 0.001, *η*^*2*^_*p*_ = 0.55), however, there were no two- or three-way interactions involving timepoint (all *p* ≥ 0.356). The main effects and interactions of age and visual feedback on adaptation index were also non-significant (all *p* ≥ 0.456).

**Figure 3.**
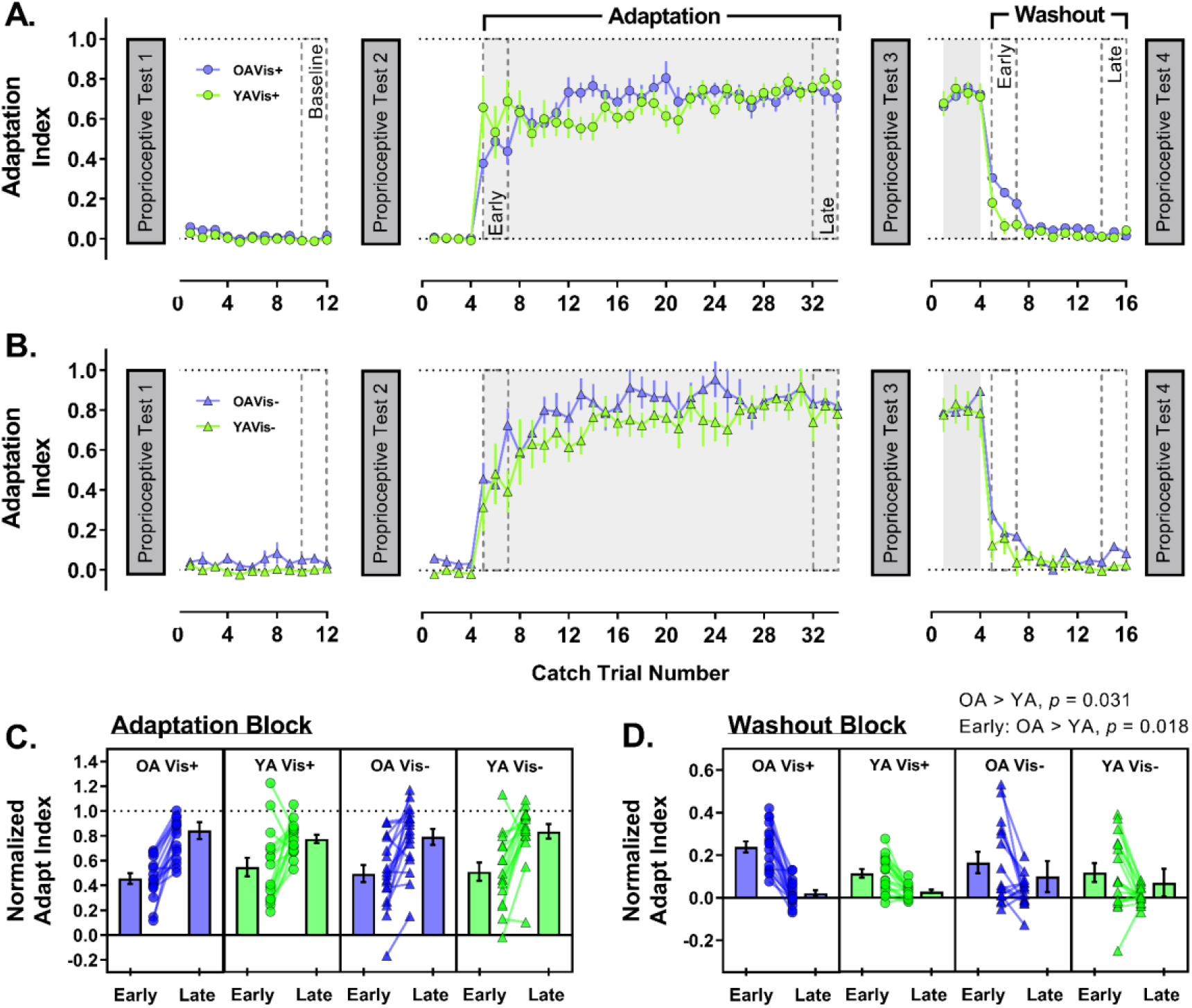
Change in adaptation index across the experiment for older (purple) and younger (green) adults in the (**A**) Vision+ condition (circles) and (**B**) Vision-condition (triangles). Lower panels show adaptation index data (normalized to baseline) for the early and late portions of the adaptation block (**C**) and the washout block (**D**). The format for each panel is as described for Figure 2.

Overall, adaptation index was reduced from early to late in the washout block (F[1, 61] = 63.23, *p* < 0.001, *η*^*2*^_*p*_ = 0.51). The timepoint x age group interaction was significant (F[1, 61] = 5.05, *p* = 0.028, *η*^*2*^_*p*_ = 0.08), where older adults (0.20 ± 0.16) maintained a larger adaptation index than younger adults (0.12 ± 0.14) early in the washout block (t[63] = 2.43, *p* = 0.018; Figure 3D). In fact, older adults (0.12 ± 0.09) maintained a larger adaptation index than younger adults (0.07 ± 0.08) across the entire washout block (F[1, 61] = 4.85, *p* = 0.031, *η*^*2*^_*p*_ = 0.07). The remaining main effects and interactions were not significant (all *p* ≥ 0.111).

#### Physical activity and adaptation

We used physical activity z-scores alongside the category variables of age and visual feedback group to predict adaptation and washout extent in a series of multiple linear regression models. We focus mainly on the strength of the physical activity coefficient as a predictor in these models, with the categorical group variables included to control for some of the interaction effects reported in the previous section where we outline a more detailed analysis of these factors. The models showed that physical activity did not predict adaptation extent (deviation error model – β = 0.12, t(62) = 0.97, *p*_adj_ = 0.505; adaptation index model – β = 0.12, t(62) = 0.89, *p*_adj_ = 0.505) or washout extent (deviation error model – β = 0.02, t(63) = 0.18, *p*_adj_ = 0.859; adaptation index model – β = −0.20, t(64) = −1.60, *p*_adj_ = 0.455), with none of the models being significant overall (all R^2^ ≤ 0.12, *p*_adj_ ≥ 0.181). This indicates that physical activity was not strongly associated with adaptation.

#### Movement kinematics

Peak velocity of movements made over course of the experiment are shown in Figure 4A (Vision+) and B (Vision-); peak velocity averaged across the adaptation block (when the force-field was switched on) is shown in Figure 4C.

**Figure 4.**
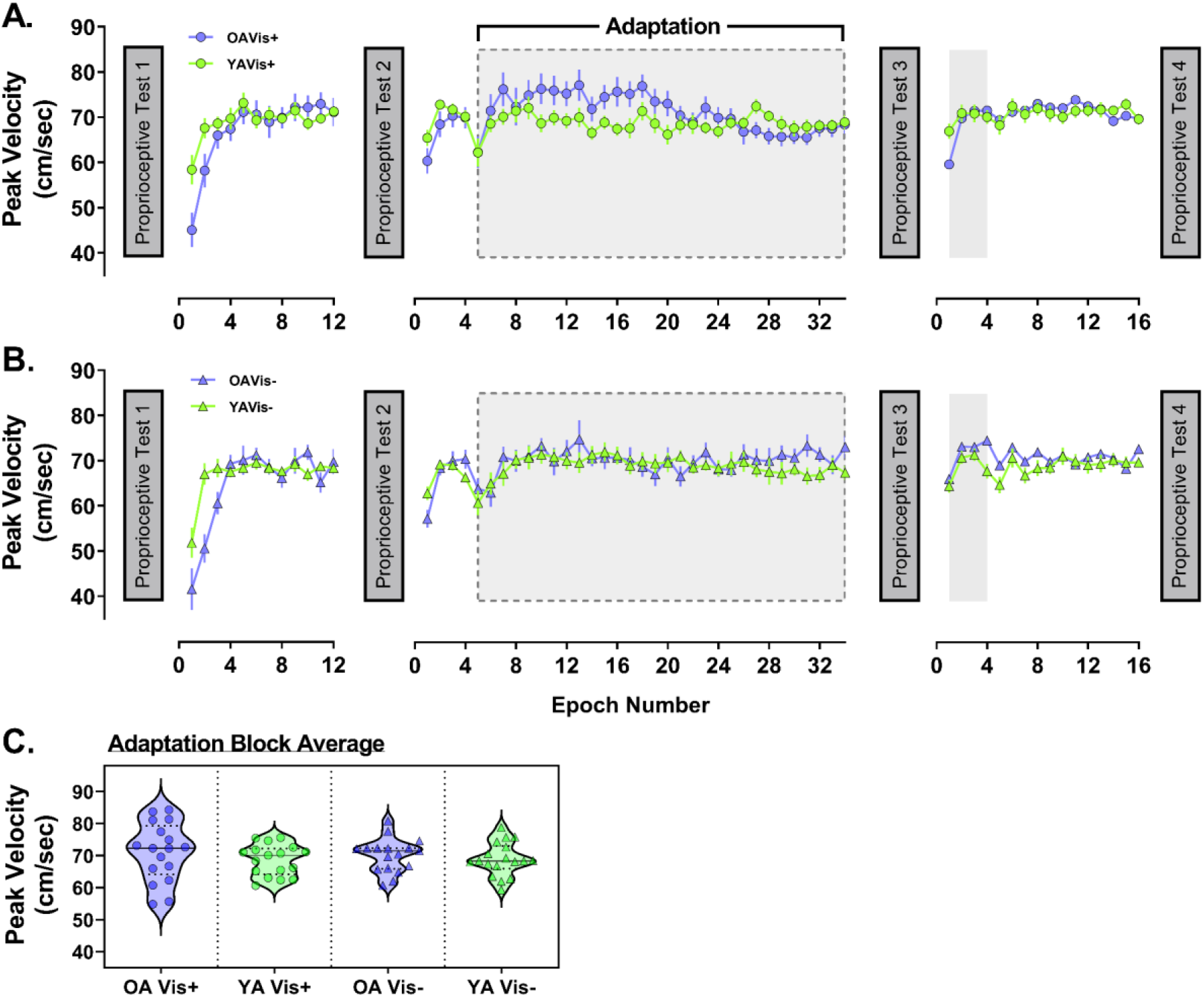
Peak velocity data shown across the experiment for older (purple) and younger (green) adults in the (**A**) Vision+ condition (circles) and (**B**) Vision-condition (triangles). For visualization, epochs of 4 trials were created for each participant and then averaged within groups to generate data points above (± 1SE), with grey shaded regions representing periods where the force-field was on. (**C**) Violin plots of peak velocity averaged across the adaptation block when the force-field was switched on (indicated by grey dashed box outline in **A** and **B**) for all four sub-groups.

A two-way (age x visual feedback group) ANOVA conducted on average peak velocity in the adaptation block showed that movement speed in the force-field was comparable between all groups (age group – F[1, 65] = 1.47, *p* = 0.230; visual feedback group – F[1, 65] = 0.06, *p* = 0.804; age x visual feedback group – F[1, 65] = 0.17, *p* = 0.686). This indicates that force exposure was similar between groups during adaptation to the velocity-dependent field. Detailed statistical analyses of baseline-normalized peak velocity, movement duration and time to peak velocity across the different phases of the experiment can be found in the Supplementary 1. As a brief summary, participants dropped peak velocity in early adaptation, but it was otherwise similar across experimental phases and between groups (Supplementary Figure S1-A). Movements became longer in duration when the perturbation was introduced, then reduced back to baseline levels and remained stable, before finally becoming shorter in duration by the end of the experiment (Supplementary Figure S1-B).

Finally, time to peak velocity (expressed as percentage of movement duration) became shorter in early adaptation, recovering slightly by late adaptation, before returning to near baseline values for the remainder of the experiment. In addition, older adults tended to have shorter time to peak velocity relative to baseline than younger adults (Supplementary Figure S1-C).

### Dynamic Proprioception

#### Baseline proprioceptive acuity

Baseline (P1) proprioceptive assessments were made before exposure to either visual feedback condition, hence a two-way (age x physical activity group) ANOVA was performed for this data (Figure 5). There was no effect of age on uncertainty range (F[1, 63] = 0.41, *p* = 0.526; Figure 5A), nor a physical activity group effect, (F[1, 63] = 0.05, *p* = 0.825) or an interaction (F[1, 63] = 0.01, *p* = 0.909). There was also no effect of age on bias (F[1, 64] = 0.55, *p* = 0.463). However, there was an effect of physical activity group (F[1, 64] = 7.64, *p* = 0.007, *η*^*2*^_*p*_ = 0.11; Figure 5B) whereby physically inactive participants had larger rightward biases overall. The age x physical activity group interaction was not significant (F[1, 64] = 0.008, *p* = 0.929). Thus, inactive participants displayed larger systematic proprioceptive errors at baseline, regardless of age.

**Figure 5.**
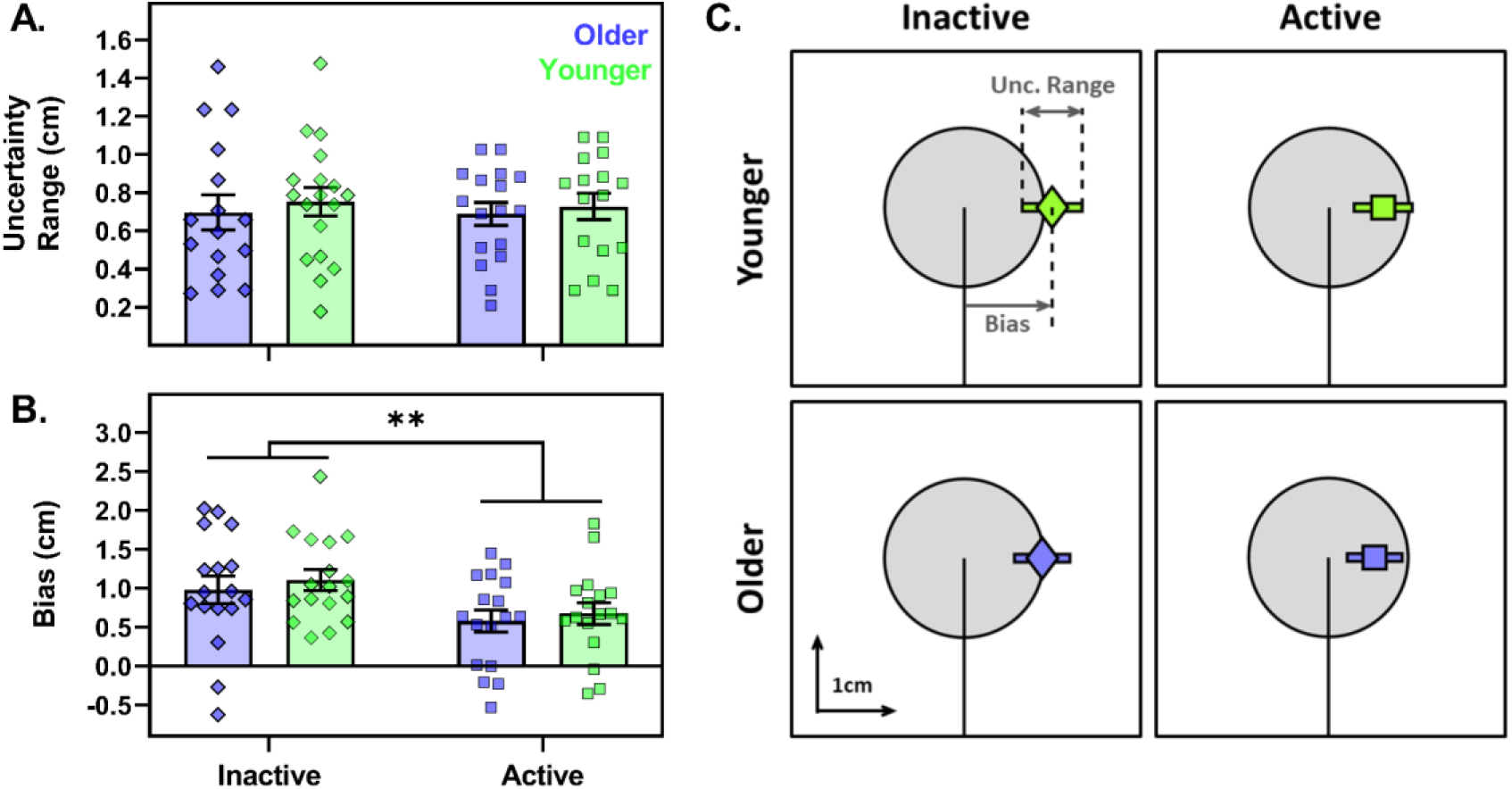
Baseline proprioceptive acuity for older (purple) and younger (green) participants grouped as physically active (squares) or inactive (diamonds). On the left-hand side, uncertainty range (**A**) and bias (**B**) are presented as group average bars (± 1SE) and individual participant symbols, where **\*\*** indicates effect of physical activity group (*p* < 0.01). **(C)** the group averaged data from panels **A** and **B**, with mean bias (diamond or square) and mean uncertainty range (bar) superimposed over the target (grey circle), at the same scale for visualisation.

#### Proprioceptive acuity during adaptation

The change in bias relative to baseline (P1) across the repeated proprioceptive assessments (P2-P4) is shown for all four groups in Figure 6, with raw baseline performance violin plots inset for reference. A three-way (age group x visual feedback x repetition) ANOVA showed no effect of repetition on the baseline-normalized bias (F[2, 122] = 1.43, *p* = 0.244), nor were there any two- or three-way interactions of repetition with either age or visual feedback group (all *p* ≥ 0.138). There was, however, a main effect of age group on normalised bias (F[1, 61] = 4.16, *p* = 0.046, *η*^*2*^_*p*_ = 0.07), where younger adults tended to have a more leftward shifted bias overall (indicated by a negative normalised bias value) than older adults. There was no main effect of visual feedback group (F[1, 61] = 0.90, *p* = 0.346) or an age group x visual feedback group interaction (F[1, 61] = 0.10, *p* = 0.756).

**Figure 6.**
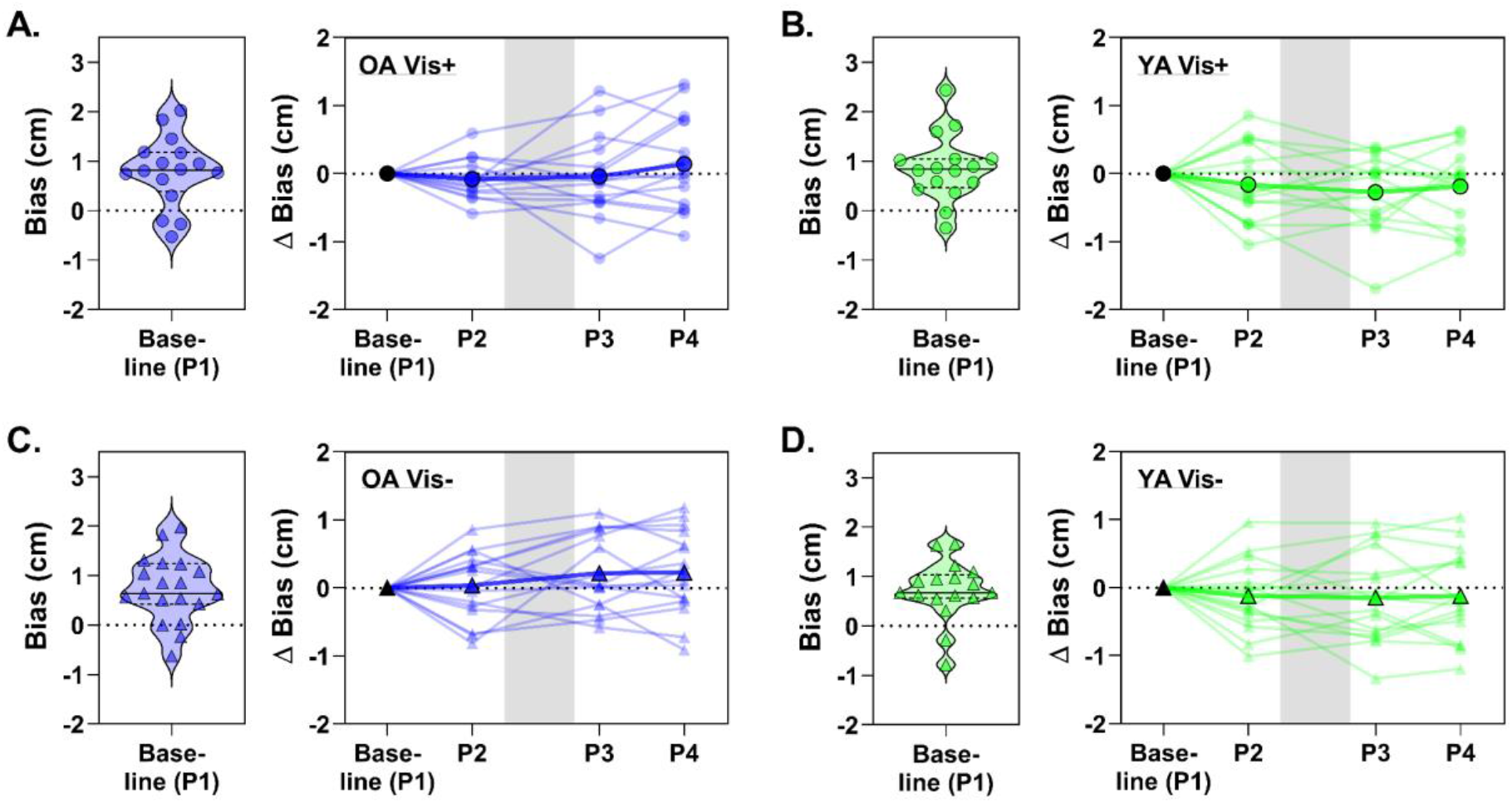
Proprioceptive bias data from all repeat assessments, with each panel showing data for one of the four groups as labelled (**A, C**: older adults; **B, D:** younger adults). Within each panel the group baseline values are shown (left, violin plot) as a visual reference for the baseline-normalized data (right, line plots). The line plots show the group average for each assessment superimposed over individual participant data, the grey shaded region represents the phase in which the adaptation block was performed.

For normalised uncertainty range, there was no main effect of repetition (F[2, 124] = 0.22, *p* = 0.804) nor any two- or three-way interactions of repetition with either age or visual feedback group (all *p* ≥ 0.311). Furthermore, there were no main effects or interactions of age or visual feedback group (all *p* ≥ 0.532).

Thus, there were no consistent changes in proprioceptive bias or uncertainty after successful adaptation to the force-field.

#### Kinematics and proprioceptive acuity

After combining the raw, non-normalised data from all four proprioceptive assessments we found that movement speed was not correlated with either uncertainty range (all |r| ≤ 0.17, *p*_*adj*_ ≥ 0.685) or absolute bias (all |r| ≤ 0.15, *p*_*adj*_ ≥ 0.717) for any of the four separate groups. This ruled out any effect of differences in proprioception due to self-selection of movement speed. Furthermore, whilst we did find that forces exerted against the channel walls were correlated with the bias for the younger vision-group, this effect did not survive adjustment for multiple comparisons (r = −0.27, *p* = 0.026, *p*_*adj*_ = 0.105) and the correlations were not significant for the remaining 3 groups (all |r| ≤ 0.20, *p*_*adj*_ ≥ 0.233). Thus, our estimates of proprioceptive acuity were largely independent of movement kinematics.

### Relationship of Proprioception and Force-Field Adaptation

#### Baseline proprioceptive acuity and adaptation

When baseline (P1) proprioceptive acuity was correlated with early and late performance in the adaptation block, there were no relationships found for either the uncertainty range (Supplementary Table S1; PVLD – all |r| ≤ 0.36, *p*_adj_ ≥ 0.556, 1.35 ≤ *BF*_*01*_ ≤ 3.24; adaptation index – all |r| ≤ 0.47, *p*_adj_ ≥ 0.462, 0.73 ≤ *BF*_*01*_ ≤ 3.32) or absolute bias (Supplementary Table S2; PVLD – all |r| ≤ 0.44, *p*_adj_ ≥ 0.598, 0.77 ≤ *BF*_*01*_ ≤ 3.32; adaptation index – all |r| ≤ 0.49, *p*_adj_ ≥ 0.211, 0.58 ≤ *BF*_*01*_ ≤ 3.34) for any of the four groups. Thus, we found little evidence to support an association between dynamic proprioceptive acuity and force-field adaptation.

#### Proprioceptive recalibration and adaptation extent

Based on previous work (Ohashi et al., 2019; Ostry et al., 2010), we expected that larger perceptual shifts in the direction of the applied perturbing force-field (indicated here by positive shift in bias) would be positively associated with greater adaptation extent. In fact, the correlations were of mixed directions and non-significant for all four groups (Supplementary Table S3; PVLD extent – all |r| ≤ 0.39, *p*_adj_ ≥ 0.505, 1.13 ≤ *BF*_*01*_ ≤ 3.34; adaptation index extent – all |r| ≤ 0.46, *p*_adj_ ≥ 0.285, 0.72 ≤ *BF*_*01*_ ≤ 3.05). Accordingly, we found no indication that proprioceptive recalibration was related to extent of adaptation.

### Spatial Working Memory

#### Effects of ageing

Memory load (6, 8, or 10 tiles) had an effect on within-search errors (WSEs; F[1.2, 70.0] = 26.86, *p* < .001, *η*^*2*^_*p*_ = 0.31; Supplementary Figure S3-A) wherein errors increased with load (post-hoc comparisons of number of tiles, all t[60] ≥ 4.01, *p*_*adj*_ < 0.001). Older adults (0.64 ± 0.60) also tended to make more WSEs than younger adults (0.38 ± 0.37) overall (F[1, 59] = 4.43, *p* = 0.040, *η*^*2*^_*p*_ = 0.07). There was no memory load x age group interaction (F[1.2, 70.0] = 1.28, *p* = 0.270).

Between-search errors (BSEs) also increased with memory load (F[1.3, 82.4] = 92.5, *p* < 0.001, *η*^*2*^_*p*_ = 0.60; all post-hoc comparisons t[64] ≥ 7.40, *p*_*adj*_ < 0.001; Supplementary Figure S3-B) and were significantly larger in older (5.12 ± 2.48) than younger adults (2.44 ± 2.10; F[1, 63] = 22.0, *p* < 0.001, *η*^*2*^_*p*_ = 0.26). Post-hoc comparisons of the significant memory load x age group interaction (F[1.3, 82.4] = 8.26, *p* = 0.002, *η*^*2*^_*p*_ = 0.12) showed that older adults made more BSEs than younger adults in the higher load conditions of 8- (t[49.5] = 4.72, *p*_*adj*_ < 0.001) and 10-tiles (t[63] = 3.54, *p*_*adj*_ = 0.001) specifically, whilst performance was similar between ages at the lowest memory load condition of 6-tiles (t[53.5] = 1.41, *p*_*adj*_ = 0.165). Thus older adults showed reduced spatial working memory capacity that became more pronounced with task complexity.

#### Working memory and adaptation/washout levels

Given that previous data has implicated spatial working memory (SWM) capacity of older adults with both adaptation (Anguera et al., 2011; Uresti-Cabrera et al., 2015) and retention (Trewartha et al., 2014) of behaviour, we correlated total search errors (sum of WSEs and BSEs across all load conditions) with early and late performance in the adaptation and washout blocks. Although we saw some evidence that younger adults with lower SWM capacity made larger movement errors in early adaptation (Vis+, r = 0.46, *p* = 0.063, *p*_*adj*_ = 0.384, *BF*_*01*_ = 0.68; Vis-, r = 0.42, *p* = 0.096, *p*_*adj*_ = 0.384, *BF*_*01*_ = 0.93), these relationships were not significant and neither were the remaining relationships with SWM in the adaptation block (Supplementary Table S4; PVLD – all remaining |r| ≤ 0.26, *p*_*adj*_ ≥ 0.730, 2.10 ≤ *BF*_*01*_ ≤ 3.33; adaptation index – all |r| ≤ 0.49, *p*_*adj*_ ≥ 0.350, 0.51 ≤ *BF*_*01*_ ≤ 3.31). or the washout block (Supplementary Table S5; PVLD – all |r| ≤ 0.33, *p*_*adj*_ ≥ 0.755, 1.52 ≤ *BF*_*01*_ ≤ 3.33; adaptation index – all |r| ≤ 0.49, *p*_*adj*_ ≥ 0.360, 0.52 ≤ *BF*_*01*_ ≤ 3.33). Thus, adaptation and SWM capacity were relatively independent in this experiment

#### Working memory and baseline proprioceptive acuity

Total search errors were not correlated with absolute baseline bias for either older (r = −0.054, *p* = 0.762, *BF*_*01*_ = 4.49) or younger adults (r = −0.182, *p* = 0.303, *BF*_*01*_ = 2.82). However, there was a trend towards an association with baseline uncertainty range in older adults (r = 0.311, *p* = 0.08, *BF*_*01*_ = 1.05) which was not seen in the younger group (r = 0.168, *p* = 0.343, *BF*_*01*_ = 3.04). This suggests that SWM capacity was largely independent of dynamic proprioceptive errors, although, there was a tendency for older participants with lower SWM capacity to have lower perceptual acuity.

## Discussion

We aimed to investigate the relationship of dynamic proprioception with adaptation to novel dynamic forces, in older and younger adults. We found that the level of adaptation was similar between the age groups, regardless of visual feedback limitations, although older adults showed increased after-effects of force production. Surprisingly, we found systematic (but not variable) proprioceptive errors were larger in physically inactive participants, regardless of age. However, there was no association between baseline proprioceptive acuity and adaptation, nor any consistent evidence of proprioceptive recalibration. Finally, despite clear age-related deficits in spatial working memory capacity there was no relation of this cognitive measure with adaptation. Taken together, we find little evidence to support a relationship between dynamic upper limb proprioception and adaptation to novel field dynamics in either older or younger adults.

This study adds to the increasing, but still limited, number of reports indicating a minimal effect of ageing on reach adaptation to novel forces (Cesqui et al., 2008; Rajeshkumar & Trewartha, 2019; Reuter et al., 2018; Trewartha et al., 2014). In addition, we demonstrate this finding holds true under conditions of limited visual feedback (providing only distance information during movement, and terminal error), conditions which might emphasize proprioceptive control. Whereas studies of force-field adaptation in the ageing population are still scarce, there is a wealth of research which has focused on visuomotor transformations in this group (usually a rotated or altered gain hand position feedback; Anguera et al., 2011; Bock, 2005; Buch et al., 2003; Contreras-Vidal et al., 2002; Hegele & Heuer, 2010; Seidler, 2006; Vandevoorde & Orban de Xivry, 2019). Although the magnitudes of age-specific deficits reported in these studies are mixed, a recurring observation is that performance is most impaired for older adults when the perturbation is salient and requires greater reliance on explicit strategies for adaptation (Buch et al., 2003; Cressman et al., 2010; Hegele & Heuer, 2010, 2013; McNay & Willingham, 1998). Converging evidence suggests this is directly related to age-dependent cognitive decline and in particular spatial working memory capacity (Anguera et al., 2011; Uresti-Cabrera et al., 2015; Vandevoorde & Orban de Xivry, 2019; Wolpe et al., 2020; for review see Seidler et al., 2010), which has been directly associated with the explicit component of adaptation (Christou et al., 2016). Despite clear evidence of reduced spatial working memory capacity in our sample of older adults, we did not find any association of this cognitive measure with adaptation.

One interpretation of these data might therefore be that force-field adaptation relies less on explicit strategies than do visuomotor rotation tasks. Specifically, this could explain why older adults appear to rely more on implicit adaptation processes (Vandevoorde & Orban de Xivry, 2019; Wolpe et al., 2020) but do not present notable deficits in force-field adaptation, which also appears to be unrelated to spatial working memory impairments. This isn’t to say that force-field adaptation relies only on implicit processes. Indeed, whilst better documented for visuomotor tasks (Benson et al., 2011; Mazzoni & Krakauer, 2006; Neville & Cressman, 2018; Taylor et al., 2014; Werner et al., 2015), recent work has sought to measure the explicit component of force-field adaptation (Schween et al., 2020) and the fast and slow processes of motor adaptation distinguished within force-field tasks (Smith et al., 2006) have been shown to be closely related to explicit and implicit learning on visuomotor tasks respectively (McDougle et al., 2015). However, the lack of any relationship with age or with age-related cognitive decline suggests explicit strategies may be reduced in force adaptation compared with visuomotor paradigms. Nevertheless, this interpretation of results was beyond the immediate scope of this study, so further research will be necessary to test this hypothesis directly.

We did not observe any consistent, direction-specific shifts in perceived hand position following adaptation, regardless of age or visual feedback conditions. Sensory perceptual recalibration with adaptation has been reported elsewhere (Cressman & Henriques, 2009; Modchalingam et al., 2019; Rossi et al., 2019; Sexton et al., 2019; Shiller et al., 2009), including proprioceptive shifts related to force-field adaptation (Haith et al., 2008; Mattar et al., 2013; Ohashi et al., 2019; Ostry et al., 2010). In the latter case, the magnitude of recalibration is typically in the range of 1-4mm (cf. Haith et al. 2008 where this is closer to 20mm), which means that factors like measurement sensitivity might contribute to difficulties in capturing these shifts empirically. Although different proprioceptive assessment methods can often provide distinct estimates of acuity (Elangovan et al., 2014; Hoseini et al., 2015), we specifically chose a task which emulates several previous studies that have reported perceptual recalibration after adaptation (Mattar et al., 2013; Ohashi et al., 2019; Ostry et al., 2010). Furthermore, we are confident in the sensitivity of our measures since we were able to detect an average 4mm bias difference between physically inactive and active participants (Figure 5B). We also note that proprioceptive recalibration has been observed despite variations of channel trajectory, movement type (passive instead of active) and staircase procedure (Ohashi et al., 2019), as well as completely different tasks altogether (Haith et al. 2008). Moreover, whilst we employed a slightly weaker force-field (15 N.m^−1^.s^−1^) than other studies reporting recalibration effects (≥ 18 N.m^−1^.s^−1^; Mattar et al., 2013; Ohashi et al., 2019; Ostry et al., 2010; Haith et al. 2008), the magnitude of movement error was in a similar range and stronger fields do not always evoke proprioceptive shifts (Sexton et al. 2018). Collectively, these results suggest that our choice of perceptual task parameters is not the primary basis for the absence of adaptation-dependent shifts in this experiment.

Consequently, factors contributing to the presence (or absence) of proprioception recalibration with force-field motor adaptation are still unclear and are likely made difficult to assess by the small scale on which recalibration occurs.

Previously, we reported that physically inactive older adults had larger systematic (but not variable) proprioceptive errors compared with a single group of younger adults (n = 20; Kitchen & Miall, 2019), which we suggested may have resulted from use-dependent intrafusal fibre sparing in advanced age that biased the tuning of sensed limb position (Bergenheim et al., 2000; Jones et al., 2001). In this experiment, an increased sample size allowed us to assign both older and younger adults into physical activity sub-groups, which revealed physically inactive participants have larger proprioceptive biases, regardless of age. Since selective loss of intrafusal fibres in low activity younger adults seems unlikely, an alternative explanation (as first suggested by Wilson et al., 2010) is that sedentary behaviour biases the experience of limb movements to a more laterally constrained region than is typically seen in naturalistic, active settings (Howard, Ingram, Körding, et al., 2009). If so, perception may be drawn towards the location of highest or most frequent sensory experience as a prior (Gritsenko et al., 2007; Körding & Wolpert, 2006), increasing systematic proprioceptive biases during periods of sensory uncertainty (as in the perceptual test). Further study with methods that permit measurements of everyday movements over extended periods of time will be necessary to test this directly.

We found that younger adults made larger lateral errors than older adults when the perturbation was first introduced (Figure 2C). Since any systematic difference in movement velocity between the age groups disappeared early in the baseline phase (Figure 4; Supplementary Figure S1-A), this raises questions as to the basis for this age-dependent difference in movement error. One explanation could be that older adults increase muscle co-contraction during movement to reduce age-dependent motor variability (Huang & Ahmed, 2014; Seidler-Dobrin et al., 1998). Increased co-contraction has been reported for older adults during force-field adaptation, but interestingly, was negatively correlated with adaptation extent (Huang & Ahmed, 2014). Unlike Huang & Ahmed (2014), we did not observe age differences in adaptation. Accordingly, it could be that our older adults reduced their initially high co-contraction over time, as observed in younger adults (Heald et al., 2018; Huang et al., 2012; Huang & Ahmed, 2014), thus minimizing maladaptive consequences on their overall performance. Ability to modulate co-contraction and subsequent limb stiffness may therefore be a factor determining adaptation in advanced age.

An unexpected finding was that older adults showed increased after-effects of force production profiles during washout (Figure 3D). This is particularly surprising given the absence of any major age effects on the adaptation phase of the experiment. Although sometimes overlooked, the washout phase or “de-adaptation” of sensorimotor adaptation tasks is considered an active process and not a simple switching-off or gradual forgetting of the behavior (Davidson & Wolpert, 2004) that can be modified with explicit knowledge and instruction (Benson et al., 2011; Mazzoni & Krakauer, 2006). Prior work with older adults has indicated reduced retention of force-field adaptation with advanced age (Rajeshkumar & Trewartha, 2019; Trewartha et al., 2014), however retention was measured in these studies using error-clamped channel trials without vision, which differs from our null-field washout phase. Elsewhere, retention differences were either equivocal or not studied in detail (Cesqui et al., 2008; Huang & Ahmed, 2014; Reuter et al., 2018). In visual perturbation tasks, after-effects are generally unaffected by age (Bock, 2005; Hegele & Heuer, 2010; see also Table 1 in Buch et al., 2003) although some studies have indicated age-specific increases in magnitude of after-effects (Fernández-Ruiz et al., 2000; Wolpe et al., 2020). This result has been suggested to depend on over-reliance on non-strategic adaptation that persists into washout (Fernández-Ruiz et al., 2000), but this notion is difficult to resolve with our data. If over-reliance on non-strategic processes was strong enough to present as increased after-effects for older adults, we would expect to see some evidence of it during the adaptation phase (as observed by Fernández-Ruiz et al., 2000), yet this wasn’t the case. Furthermore, the absence of an association with working memory at least suggests that an age-dependent imbalance in the use of cognitive strategies was unlikely to have contributed to the effect. Accordingly, further study on factors which influence this aspect of performance in advanced age are necessary to better understand the basis of this novel finding.

In conclusion, we found minimal age-dependent differences in force-field adaptation and only limited evidence for proprioceptive recalibration with adaptation, regardless of visual feedback extent. We did find differences in proprioceptive acuity between high and low physical activity individuals, but this was across the age range. Dynamic proprioceptive acuity deficits at the non-clinical level (with healthy ageing or physical inactivity) do not, therefore, appear to influence force-field adaptation. Age-dependent cognitive decline has been closely linked with adaptation deficits in visuomotor tasks, but we found no association between spatial working memory capacity and adaptation performance. Hence, we suggest that force-field adaptation may be sufficiently weighted towards implicit processes to abolish these age-dependent performance impairments.

## Acknowledgements

This work was funded by the MRC-ARUK Centre for Musculoskeletal Ageing Research (CMAR) with support from the Wellcome Grant WT087554 and NIH/NIDCD R01DC014510.

