## Supplementary Material for "Adaptation of reach action to a novel force-field is not predicted by acuity of dynamic proprioception in either older or younger adults"

### 1. Extended Analysis of Force-Field Adaptation Kinematics

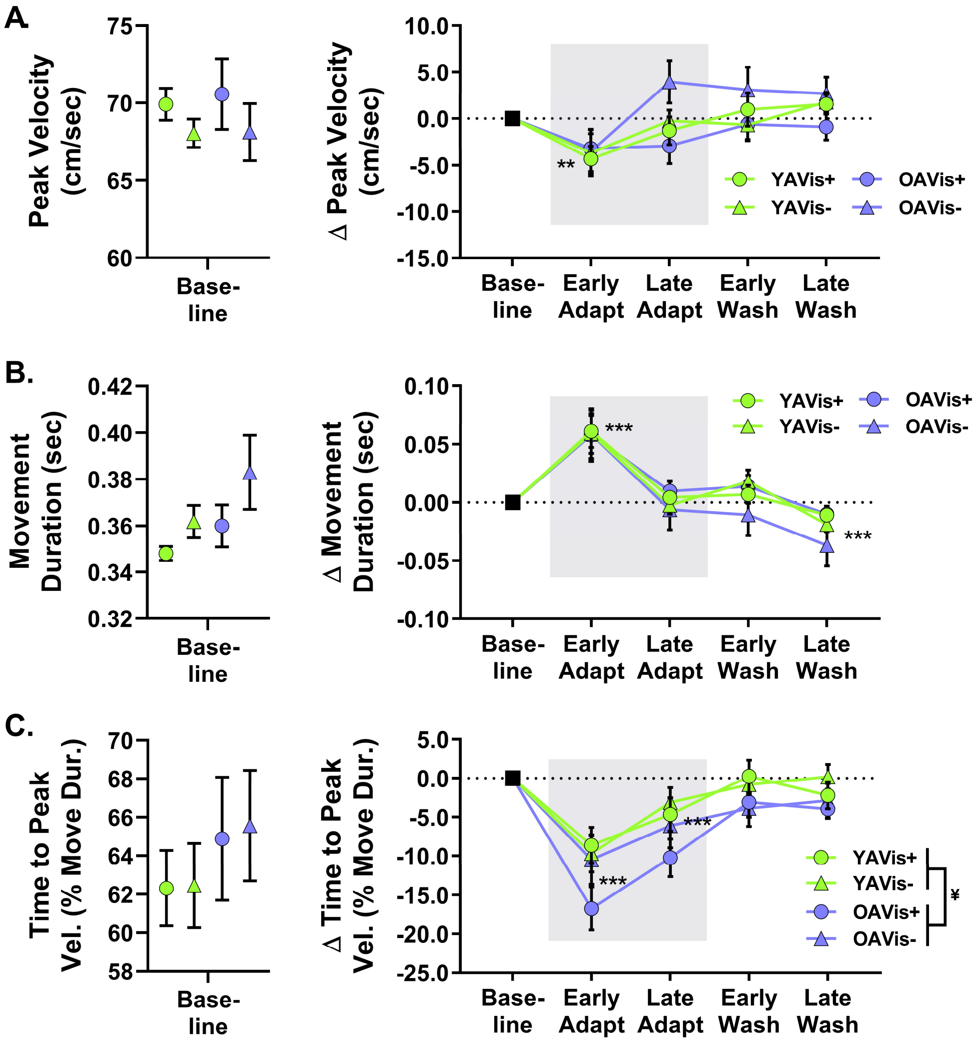


**Figure S1**. – Group average (± 1SE) kinematic data for the force-field adaptation task across different phases of the experiment. All data is normalized to baseline (late null) performance which is shown in the left of each panel. **A.** shows the peak velocity data, **B.** shows the movement duration and **C.** shows the time to peak velocity represented as a percentage of total movement duration. Significant differences between phases is shown as ** ($p_{adj}$ < 0.01), *** ($p_{adj}$ < 0.001) and effect of age group indicated by ¥ ($p$ < 0.05).

**Normalized peak velocity** – There was an effect of phase on normalised peak velocity ($F[2.5, 152.5] = 10.0$, $p < 0.001$, $\eta^2_p = 0.14$) with follow-up comparisons indicating that peak velocity dropped in early adaptation compared to all other phases (all $t[64] \geq 3.21$, $p_{adj} \leq 0.005$ – Figure S1-A). There were no two- or three-way interactions of phase (all $p > 0.262$), as well as no effects or interactions of vision and age group on normalised peak velocity (all $p > 0.180$).

**Normalized movement duration** – There was also an effect of phase on normalised movement duration ($F[1.9, 117.1] = 35.6$, $p < 0.001$, $\eta^2_p = 0.36$) where movement duration increased in early adaptation compared to all other phases (all $t[66] \geq 5.40$, $p_{adj} < 0.001$), but was similar from late adaptation to early washout ($t[66] = -1.0$, $p_{adj} = 0.319$), and then finally reduced in late washout compared to all other phases (all $t[66] \geq 3.85$, $p_{adj} < 0.001$ – Figure S1-B). The two- and three-way interactions of phase were not significant (all $p > 0.588$), and there were no effects or interactions of vis and age group (all $p > 0.318$).

**Normalized time to peak velocity** – There was an effect of phase on normalised time to peak velocity (TPV – $F[2.4, 152.8] = 28.6$, $p < 0.001$, $\eta^2_p = 0.31$), where TPV dropped to its shortest in early adaptation ($t[67] \geq 3.85$, $p_{adj} < 0.001$ compared to all other phases), recovered towards baseline by late adaptation (all $t[67] \geq 3.85$, $p_{adj} < 0.001$ compared to other phases) and then remained stable at near baseline levels during washout (early vs late washout; $t[67] = 0.38$, $p_{adj} = 0.703$ – Figure S1-C). However, there were no two- or three-way phase interactions on normalised TPV (all $p > 0.274$). There was no between-subjects effect of vis group on normalised TPV ($F[1, 64] = 0.87$, $p = 0.354$), however there was an effect of age group on TPV ($F[1, 64] = 4.6$, $p = 0.036$, $\eta^2_p = 0.07$) where overall, older adults reduced their TPV relative to baseline (and hence reached peak velocity earlier in their movements) more than younger adults. The vis group x age group interaction was not significant ($F[1, 64] = 0.43$, $p = 0.511$).

### 2. Proprioception and Force-Field Adaptation Correlation Tables

**BL Uncertainty Range vs. Adaptation**

**PVLD**

| Group | | Early | | | | Late | | | |
|---|---|---|---|---|---|---|---|---|---|
| | | *r* | *p* | $p_{fdr}$ | $BF_{01}$ | *r* | *p* | $p_{fdr}$ | $BF_{01}$ |
| Older | Vis+ | 0.341 | 0.196 | 0.556 | 1.497 | 0.113 | 0.689 | 0.982 | 2.918 |
| | Vis- | -0.006 | 0.982 | 0.982 | 3.241 | -0.071 | 0.785 | 0.982 | 3.226 |
| Younger | Vis+ | -0.355 | 0.162 | 0.556 | 1.346 | -0.042 | 0.879 | 0.982 | 3.207 |
| | Vis- | -0.086 | 0.742 | 0.982 | 3.174 | -0.332 | 0.209 | 0.556 | 1.562 |

**Adaptation Index**

| Group | | Early | | | | Late | | | |
|---|---|---|---|---|---|---|---|---|---|
| | | *r* | *p* | $p_{fdr}$ | $BF_{01}$ | *r* | *p* | $p_{fdr}$ | $BF_{01}$ |
| Older | Vis+ | -0.363 | 0.167 | 0.462 | 1.341 | -0.473 | 0.075 | 0.462 | 0.734 |
| | Vis- | 0.026 | 0.920 | 0.920 | 3.323 | 0.217 | 0.404 | 0.462 | 2.416 |
| Younger | Vis+ | 0.266 | 0.319 | 0.462 | 2.070 | 0.296 | 0.249 | 0.462 | 1.802 |
| | Vis- | 0.230 | 0.375 | 0.462 | 2.317 | 0.322 | 0.224 | 0.462 | 1.639 |

**Table S1.** – Group correlations between baseline (BL) uncertainty range and performance early and late adaptation block, as measured separately by peak-velocity lateral deviation (PVLD) and adaptation index. Correlation coefficients (*r*) are provided alongside raw (*p*) and false discovery rate-adjusted ($p_{fdr}$) *p*-values. Bayes factor for correlation indicates increasing likelihood for null as $BF_{01}$ increases from 1 and increasing likelihood for alternative as $BF_{01}$ decreases from 1 (see main text for further details).

**BL Absolute Bias vs. Adaptation**

**PVLD**

| **Group** | | **Early** | | | | **Late** | | | |
|---|---|---|---|---|---|---|---|---|---|
| | | *r* | *p* | $p_{fdr}$ | $BF_{01}$ | *r* | *p* | $p_{fdr}$ | $BF_{01}$ |
| Older | Vis+ | -0.131 | 0.616 | 0.821 | 2.970 | -0.202 | 0.454 | 0.726 | 2.500 |
| | Vis- | 0.299 | 0.261 | 0.726 | 1.808 | 0.443 | 0.075 | 0.598 | 0.767 |
| Younger | Vis+ | -0.199 | 0.444 | 0.726 | 2.545 | -0.044 | 0.871 | 0.918 | 3.202 |
| | Vis- | 0.277 | 0.282 | 0.726 | 1.951 | 0.027 | 0.918 | 0.918 | 3.322 |

**Adaptation Index**

| **Group** | | **Early** | | | | **Late** | | | |
|---|---|---|---|---|---|---|---|---|---|
| | | *r* | *p* | $p_{fdr}$ | $BF_{01}$ | *r* | *p* | $p_{fdr}$ | $BF_{01}$ |
| Older | Vis+ | -0.017 | 0.949 | 0.999 | 3.332 | -0.191 | 0.478 | 0.765 | 2.568 |
| | Vis- | -0.272 | 0.290 | 0.581 | 1.988 | -0.478 | 0.052 | 0.211 | 0.586 |
| Younger | Vis+ | 0.492 | 0.053 | 0.211 | 0.575 | < 0.001 | 0.999 | 0.999 | 3.336 |
| | Vis- | -0.321 | 0.209 | 0.556 | 1.604 | -0.146 | 0.576 | 0.767 | 2.886 |

**Table S2.** – Group correlations between baseline (BL) absolute bias and performance early and late adaptation block, measured separately by peak-velocity lateral deviation (PVLD) and adaptation index. Correlation coefficients (r) are provided alongside raw (*p*) and false discovery rate-adjusted ($p_{fdr}$) *p*-values. Bayes factor for correlation indicates increasing likelihood for null as $BF_{01}$ increases from 1 and increasing likelihood for alternative as $BF_{01}$ decreases from 1 (see main text for further details).

**Adaptation Extent vs. Bias Shift**

| **Group** | | **PVLD** | | | |
|---|---|---|---|---|---|
| | | *r* | *p* | $p_{fdr}$ | $BF_{01}$ |
| Older | Vis+ | 0.255 | 0.340 | 0.679 | 2.127 |
| | Vis- | 0.386 | 0.126 | 0.505 | 1.131 |
| Younger | Vis+ | 0.009 | 0.973 | 0.998 | 3.240 |
| | Vis- | 0.001 | 0.998 | 0.998 | 3.339 |

| **Group** | | **Adaptation Index** | | | |
|---|---|---|---|---|---|
| | | *r* | *p* | $p_{fdr}$ | $BF_{01}$ |
| Older | Vis+ | -0.354 | 0.163 | 0.326 | 1.355 |
| | Vis- | 0.462 | 0.071 | 0.285 | 0.724 |
| Younger | Vis+ | -0.142 | 0.600 | 0.661 | 2.854 |
| | Vis- | 0.115 | 0.661 | 0.661 | 3.054 |

**Table S3.** – Group correlations between adaptation extent and proprioceptive recalibration (bias shift) across the adaptation block. Adaptation extent is measured separately by peak-velocity lateral deviation (PVLD) and adaptation index. Correlation coefficients (r) are provided alongside raw (*p*) and false discovery rate-adjusted ($p_{fdr}$) *p*-values. Bayes factor for correlation indicates increasing likelihood for null as $BF_{01}$ increases from 1 and increasing likelihood for alternative as $BF_{01}$ decreases from 1 (see main text for further details). In all cases, positive correlations indicate greater adaptation was associated with greater shifts in the direction of the perturbing force-field.

### 3. Age Effects and Correlations of Spatial Working Memory Capacity

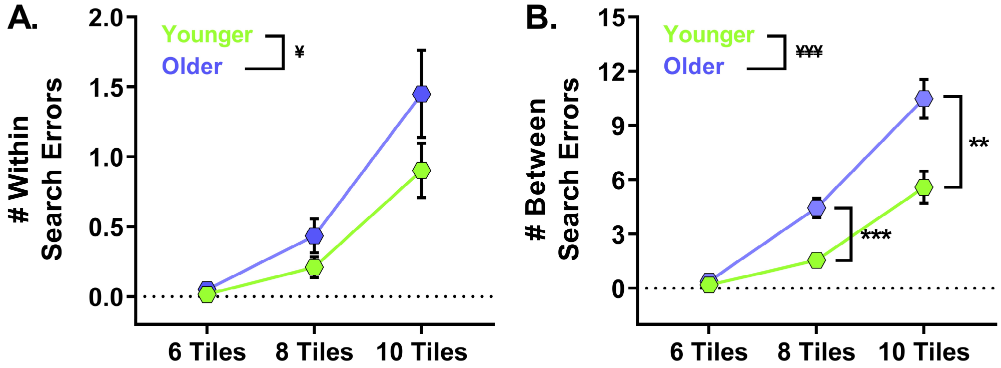


**Figure S2.** – Group average (± 1SE) search errors for older (purple) and younger (green) adults in the spatial working memory task, where task difficulty increased with number of tiles. **A.** Number of within search errors, where main effect of age is indicated by [¥] ($p < 0.05$). **B.** Between search errors, where main effect of age is indicated by [¥¥¥] ($p < 0.001$) and significant post-hoc comparisons of older and younger adults are indicated by ** ($p_{adj} < 0.01$) and *** ($p_{adj} < 0.001$).

**Total SWM Search Errors vs. Adaptation**

**PVLD**

| Group | | Early | | | | Late | | | |
|---|---|---|---|---|---|---|---|---|---|
| | | *r* | *p* | $p_{fdr}$ | $BF_{01}$ | *r* | *p* | $p_{fdr}$ | $BF_{01}$ |
| Older | Vis+ | -0.094 | 0.721 | 0.940 | 3.146 | 0.201 | 0.456 | 0.730 | 2.509 |
| | Vis- | -0.232 | 0.386 | 0.730 | 2.292 | 0.032 | 0.902 | 0.940 | 3.315 |
| Younger | Vis+ | 0.460 | 0.063 | 0.384 | 0.678 | 0.260 | 0.331 | 0.730 | 2.095 |
| | Vis- | 0.417 | 0.096 | 0.384 | 0.925 | 0.020 | 0.940 | 0.940 | 3.330 |

**Adaptation Index**

| Group | | Early | | | | Late | | | |
|---|---|---|---|---|---|---|---|---|---|
| | | *r* | *p* | $p_{fdr}$ | $BF_{01}$ | *r* | *p* | $p_{fdr}$ | $BF_{01}$ |
| Older | Vis+ | -0.038 | 0.885 | 0.885 | 3.306 | -0.074 | 0.784 | 0.885 | 3.131 |
| | Vis- | 0.076 | 0.772 | 0.885 | 3.211 | 0.211 | 0.417 | 0.885 | 2.458 |
| Younger | Vis+ | -0.345 | 0.191 | 0.763 | 1.470 | -0.494[¥] | 0.044 | 0.350 | 0.509 |
| | Vis- | -0.102 | 0.696 | 0.885 | 3.111 | 0.159 | 0.542 | 0.885 | 2.806 |

**Table S4.** – Group correlations between total search errors in the spatial working memory (SWM) task and early and late performance in the adaptation block, measured separately by peak-velocity lateral deviation (PVLD) and adaptation index. Correlation coefficients (r) are provided alongside raw (*p*) and false discovery rate-adjusted ($p_{fdr}$) *p*-values. Significant correlations that did not survive corrections for multiple comparisons indicated by [¥]. Bayes factor for correlation indicates increasing likelihood for null as $BF_{01}$ increases from 1 and increasing likelihood for alternative as $BF_{01}$ decreases from 1 (see main text for further details).

**Total SWM Search Errors vs. Washout**

**PVLD**

| Group | | Early | | | | Late | | | |
|---|---|---|---|---|---|---|---|---|---|
| | | *r* | *p* | $p_{fdr}$ | $BF_{01}$ | *r* | *p* | $p_{fdr}$ | $BF_{01}$ |
| Older | Vis+ | -0.196 | 0.452 | 0.755 | 2.567 | -0.265 | 0.303 | 0.755 | 2.042 |
| | Vis- | -0.066 | 0.802 | 0.917 | 3.243 | -0.155 | 0.566 | 0.755 | 2.785 |
| Younger | Vis+ | 0.016 | 0.950 | 0.950 | 3.332 | -0.180 | 0.489 | 0.755 | 2.672 |
| | Vis- | -0.333 | 0.192 | 0.755 | 1.516 | -0.170 | 0.500 | 0.755 | 2.778 |

**Adaptation Index**

| Group | | Early | | | | Late | | | |
|---|---|---|---|---|---|---|---|---|---|
| | | *r* | *p* | $p_{fdr}$ | $BF_{01}$ | *r* | *p* | $p_{fdr}$ | $BF_{01}$ |
| Older | Vis+ | -0.132 | 0.614 | 0.702 | 2.966 | 0.274 | 0.286 | 0.573 | 1.971 |
| | Vis- | 0.366 | 0.148 | 0.573 | 1.269 | 0.170 | 0.544 | 0.702 | 2.651 |
| Younger | Vis+ | -0.283 | 0.271 | 0.573 | 1.903 | 0.014 | 0.957 | 0.957 | 3.334 |
| | Vis- | 0.492[¥] | 0.045 | 0.360 | 0.520 | 0.222 | 0.392 | 0.627 | 2.375 |

**Table S5.** – Group correlations between total search errors in the spatial working memory (SWM) task and early and late performance in the washout block, measured separately by peak-velocity lateral deviation (PVLD) and adaptation index. Correlation coefficients (r) are provided alongside raw (*p*) and false discovery rate-adjusted ($p_{fdr}$) *p*-values. Significant correlations that did not survive corrections for multiple comparisons indicated by ¥. Bayes factor for correlation indicates increasing likelihood for null as $BF_{01}$ increases from 1 and increasing likelihood for alternative as $BF_{01}$ decreases from 1 (see main text for further details).
